# Mechanisms of Pathogenicity of Hypertrophic Cardiomyopathy-Associated Troponin T (TNNT2) Variant R278C^+/-^ During Development

**DOI:** 10.1101/2023.06.06.542948

**Authors:** Sanam Shafaattalab, Alison Y Li, Farah Jayousi, Yasaman Maaref, Saif Dababneh, Homa Hamledari, Dina Hosseini Baygi, Tiffany Barszczewski, Balwinder Ruprai, Shayan Jannati, Raghu Nagalingam, Austin M Cool, Paulina Langa, Mu Chiao, Thomas Roston, R John Solaro, Shubhayan Sanatani, Christopher Toepfer, Steffen Lindert, Philipp Lange, Glen F Tibbits

**Author notes:** These authors contributed equally to this study (Sanam Shafaattalab, Alison Y Li, Farah Jayousi, Yasaman Maaref, Saif Dababneh). Correspondence: Glen F Tibbits, PhD, Cellular and Regenerative Medicine Centre BC Children’s Hospital Research Institute 938 West 28th Avenue, Vancouver, BC, V5Z 4H4 Canada.

## Abstract

Hypertrophic cardiomyopathy (HCM) is one of the most common heritable cardiovascular diseases and variants of *TNNT2* (cardiac troponin T) are linked to increased risk of sudden cardiac arrest despite causing limited hypertrophy. In this study, a *TNNT2* variant, R278C^+/-^, was generated in both human cardiac recombinant/reconstituted thin filaments (hcRTF) and human-induced pluripotent stem cells (hiPSCs) to investigate the mechanisms by which the R278C^+/-^ variant affects cardiomyocytes at the proteomic and functional levels. The results of proteomics analysis showed a significant upregulation of markers of cardiac hypertrophy and remodeling in R278C^+/-^ vs. the isogenic control. Functional measurements showed that R278C^+/-^ variant enhances the myofilament sensitivity to Ca^2+^, increases the kinetics of contraction, and causes arrhythmia at frequencies >75 bpm. This study uniquely shows the profound impact of the *TNNT2* R278C^+/-^ variant on the cardiomyocyte proteomic profile, cardiac electrical and contractile function in the early stages of cardiac development.

**Translational Perspective:** Hypertrophic cardiomyopathy (HCM) is the leading known cause of sudden cardiac arrest in the young. Thin-variant associated HCM variants make up to 15% of familial HCM yet their molecular mechanisms remain less clear relative to thick filament variants. Here, we employ computational modeling, human cardiac recombinant/reconstituted thin filaments (hcRTF), and hiPSC-CMs to study the thin filament *TNNT2* R278C^+/-^ variant, revealing its extensive pathogenicity and potential mechanisms by which it can lead to HCM and sudden death. Mavacamten, the recently FDA-approved treatment, was effective in alleviating contractile dysfunction in *TNNT2* R278C^+/-^ hiPSC-CMs, positing it as a potential therapy for thin filament HCM.

**Graphical Abstract:** 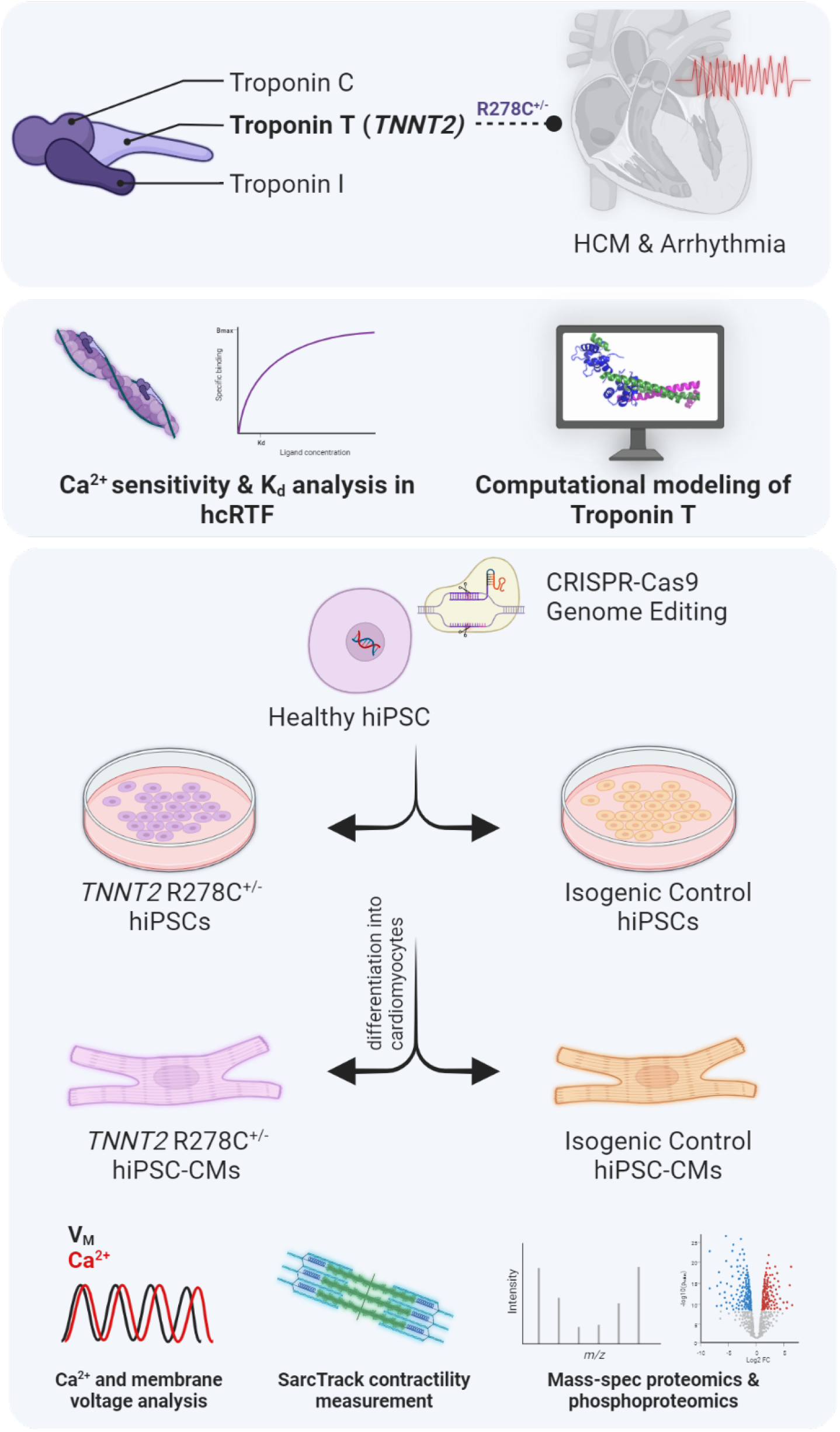

## 1. INTRODUCTION

Hypertrophic cardiomyopathy (HCM) is one of the most common inherited heart diseases and is associated with >1500 variants located in ∼20 genes with the majority encoding for sarcomeric and Z-disc proteins^1^. HCM is an autosomal dominant inherited disorder that is characterized by asymmetrical left ventricular hypertrophy, enhanced contractility, impaired relaxation, and myofibrillar disarray, which clinically can manifest sudden cardiac arrest (SCA)^2, 3^. The most common variants are in sarcomeric genes encoding the thick filament proteins cardiac myosin-binding protein C (*MYBPC3*) and β-myosin heavy chain (*MYH7*). However, variants in thin filament encoding proteins, such as *TNNT2*, make up 5-15% of HCM cases yet their pathogenic mechanisms remain relatively less clear^4, 5^. Moreover, whether mavacamten, the allosteric myosin ATPase inhibitor recently approved by the FDA to treat obstructive HCM, would be effective in *TNNT2* variant carriers has yet to be studied.

Variants in *TNNT2* represent up to 50% of all thin filament HCM cases, with 400 unique TNNT2 variants have been identified in HCM patients^5, 6^. In particular, the *TNNT2* R278C variant has been reported in over 50 unrelated patients with a clinical diagnosis of HCM ^7, 8^, and is enriched in the HCM population compared to the general population^9^. Patients carrying this variant tend to present with relatively later-onset and milder hypertrophy compared to patients with *MYH7* and *MYBCP3* associated HCM but may have a higher risk of sudden cardiac death at young ages^10–15^. However, conflicting results from several mechanistic studies in patient myectomy tissue and animal models and the presence of multiple potentially disease-associated variants in some affected individuals have raised uncertainty as to the pathogenicity of the *TNNT2* R278C variant.

Genetic variants associated with HCM can result in dramatic defects in the biophysical dynamics of Ca^2+^ handling and contractility^3, 16, 17^, which signal significant alterations in the transcriptome, proteome, and function. Recent studies have used hiPSC-CMs based disease models to investigate the underlying causes and consequences of HCM-linked variants^18, 19^. We have previously shown that the HCM-associated *TNNT2* I79N^+/-^ variant can enhance myofilament Ca^2+^ sensitivity by decreasing the Ca^2+^ off rate constant, alter the action potential morphology, and modify contractile kinetics in human iPSC-CMs^20^.

Given the conflicting evidence of pathogenicity and the paucity of studies on the *TNNT2* R278C^+/-^ variant in a human-based model, we sought to investigate R278C^+/-^ using human iPSC-CMs and human cardiac recombinant/reconstituted thin filaments (hcRTF). There is compelling evidence that the arginine at residue 278 is highly conserved over four million years of evolution (Figure S1), with several variants at this residue considered pathogenic^10, 21^. Moreover, our molecular mechanics potential energy minimization calculations strongly suggest the R278C^+/-^ variant alters the interaction of *TNNT2* with tropomyosin (Tm) (Figure 1). Using hcRTF, we found that R278C^+/-^ increased myofilament Ca^2+^ sensitivity. We then generated heterozygous *TNNT2* R278C^+/-^ hiPSC-CMs and isogenic controls and conducted electrophysiological, contractile, and proteomic analyses. R278C^+/-^ hiPSC-CMs showed various proteomic changes consistent with HCM such as myofibrillar disarray, hypercontractility, and a fetal-like metabolic switch. Using the MATLAB based algorithm SarcTrack^22^, we found sarcomere shortening, contraction, and relaxation kinetics to be disrupted in R278C^+/-^ hiPSC-CMs, which were alleviated in part by mavacamten. Lastly, optical mapping showed a higher propensity for arrhythmias induced by the R278C^+/-^ variant in hiPSC-CMs. This unique approach in studying the early impact of the *TNNT2* HCM-related variant R278C^+/-^ on cardiomyocytes has generated a magnitude of novel data that could be impactful on understanding this disease.

**Figure 1.**
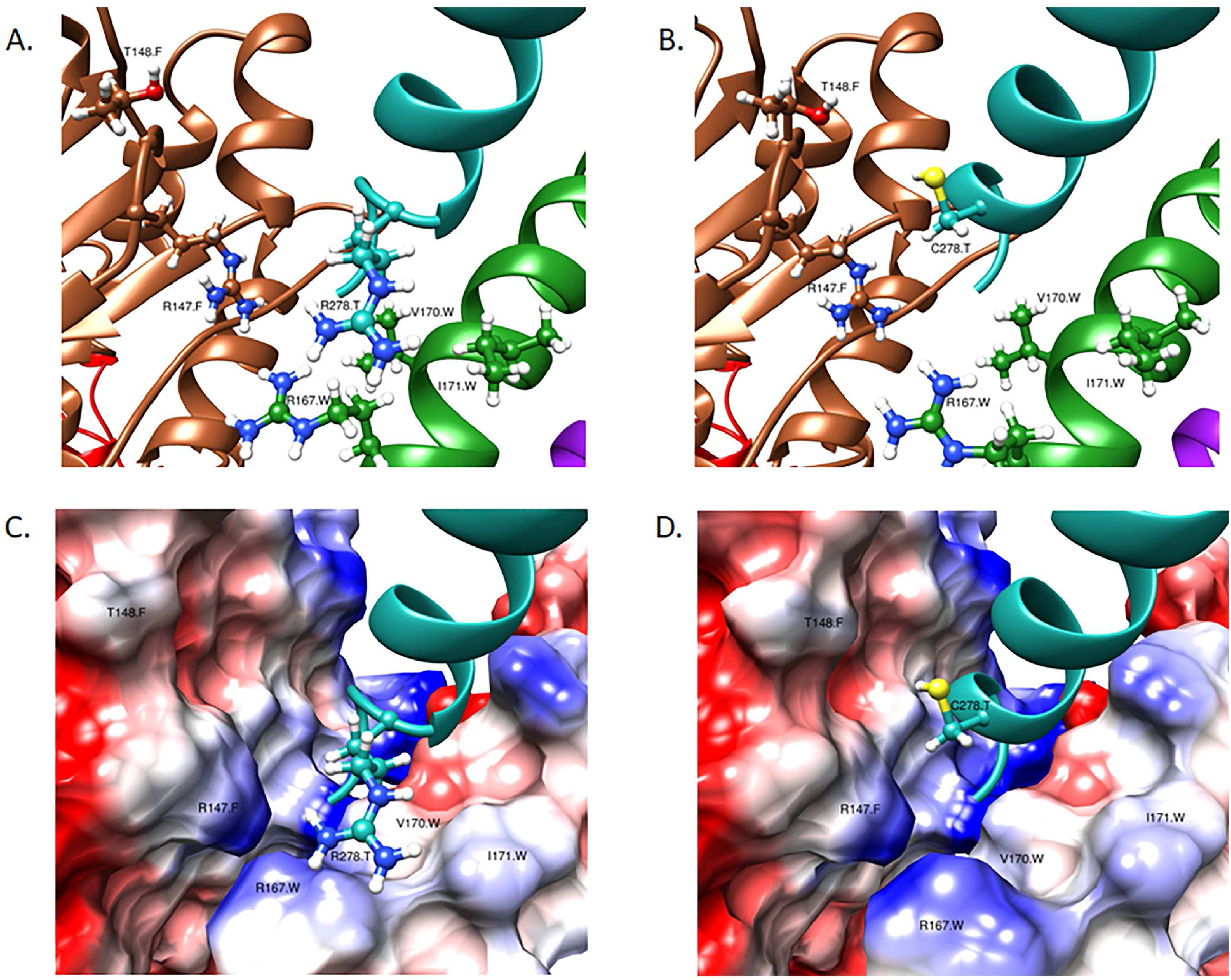
Computational modelling and potential energy minimization. Panels A. and B. show troponin T (cyan), actin chain F (brown), tropomyosin chain W (green), and tropomyosin chain X (purple). Panel A. shows the structure of the complex with WT troponin T (with residue R278). Panel B. includes the predicted structure of the complex with the genetic variant R278C^+/-^. Residues with side chain atoms displayed correspond to the five closest residues that the R278 CG (γ carbon) atom and C278 SG (γ sulfur) atom have in common. Panels C. and D. show troponin T (cyan) and the surfaces of actin chain F and tropomyosin chain W. The surfaces of these chains are colored with respect to their Coulombic surface potential, where blue indicates positive residues, red indicates negative residues, and white corresponds to neutral residues. Panel C. shows the structure of the complex with R278, and Panel D. shows the modelled structure of the Tn complex that harbors the mutant residue C278. Labels shown are in the format: one letter residue code, residue number, chain ID.

## 2. METHODS

### 2.1 Computational Modelling and Potential Energy Minimization

To obtain a model of the thin filament complex, the PDB structures 6KN7 (Ca^2+^ free) and 6KN8 (Ca^2+^ bound) were used^23^. Each of these structures contains two troponin complexes, with cardiac troponin T (cTnT) being designated as chains “a” and “T”, respectively. Both cTnT chains have contacting actin and tropomyosin subunits, in which chain a is close to chains G (actin) and P (tropomyosin) and chain T is close to chains F (actin) and W (tropomyosin). None of the four TnT structures (6KN7a, 6KN7T, 6KN8a, and 6KN8T) contains residue R278, with each cTnT structure being truncated at residue Q272.

Rosetta was utilized to build residues 273-281, using the sequence: KVSKTRGKA. Each of the four TnT chains (6KN7a, 6KN7T, 6KN8a, and 6KN8T) was removed from the thin filament complex and 1,000 models were built of the existing structure plus residues 273-281 using the loop modeling application of Rosetta^24, 25^. This was done twice, the first designated all the residues to be built as a helical structure, the second designated all the residues to be built as an unstructured loop. In both cases, Rosetta built residues 273-281 as a helical structure. The top scoring model was then placed into its original PDB structure to generate a thin filament model that contained the cTnT R278 residue.

Four more models were then created by introducing the R278C^+/-^ variant into the models produced by Rosetta. Each of the eight models of the thin filament complex were then solvated with explicit TIP3W water molecules and NaCl counterions were added to neutralize the entire system and establish a final salt concentration of 150 mM. The resulting models were subjected to two sequential 10,000 potential energy minimization steps using Nanoscale Molecular Dynamics (NAMD) in the Charmm36 force field. This was done to optimize the positions of the solvent molecules around the protein and to subsequently relax the positions of residue side chain atoms. To analyze the models, residues that were within a certain cutoff of the gamma carbon (CG) atom of the R278 residue of the native structure and of the gamma sulfur (SG) atom of the C278 residue in the mutated models, were recorded.

### 2.2 Generation of human cardiac reconstituted thin filaments (hcRTF)

Recombinant human adult troponin subunits were used to generate the cardiac troponin complexes (cTn). Recombinant human α-Tm produced in E. coli does not undergo an acetylation process and the unacetylated initial Met binds actin poorly and does not polymerize. In this study, the recombinant αTm included an MAS peptide (Met-Ala-Ser) added to the N-terminus to mimic the acetylation process and ensure actin binding^26^. The purification of each protein was verified by gel electrophoresis. Reconstituted thin filaments were then generated by mixing each cTn complex with human αMAS TPM1 as well as native actin extracted from rabbit skeletal muscle which has a 99.2% sequence identity with human cardiac actin^27^. The stopped flow assay used for testing the functional validity of the hcRTF in this study has been described by Davis et *al.* ^28^.

### 2.3 Determination of hcRTF Ca^2+^ kinetics

The Ca^2+^ off rate constants (k_off_ (Ca^2+^)) from each hcRTF were characterized using a Chirascan stopped-flow apparatus (Applied Photophysics, Surrey, UK) with a dead time of 1.1 ms at 15.0 ± 0.1°C. 2-(4’-(iodoacetoamido)anilino) naphthalene-6-sulfonic acid (IAANS) fluorophore was attached to residue C53 of TnC, was excited at 300 nm and monitored through a 510 nm broad band-pass interference filter (Semrock, Rochester, NY) as described by Davis *et al.*^28^. Each trace was obtained as the solution containing hcRTF saturated with 200 μM CaCl2 was rapidly mixed with 10 mM EGTA solution in the stopped-flow buffer containing (in mM): 10 MOPS, 150 KCl, 3 MgCl2, 1 DTT, pH 7.0. Each k_off_(Ca) value represents an average of at least five traces fit with a single exponential equation and repeated more than 15 times. Two means were considered to be significantly different when p < 0.05.

All steady-state fluorescence measurements were performed using a Cary Eclipse fluorescence spectrophotometer (Agilent, Santa Clara, CA) at 15 ± 0.1°C. In brief, the IAANS fluorophore was excited at 330 nm and monitored at the peak of the emission at 450 nm. Microliter amounts of CaCl_2_ were added to 2 mL of each labeled hcRTF in a titration buffer containing (in mM): 200 MOPS, 150 KCl, 3 MgCl2, and 1 DTT at pH 7.0. At least five to eight datasets were collected for each construct and the data were fit with a sigmoidal fit (based on the Hill equation) in Origin 8.5 (Microcal Software, Northampton, MA). The Ca^2+^ sensitivity for each hcRTF was represented by the dissociation constant K_d_ (± SEM) defined as the Ca^2+^ concentration at the half-maximal fluorescence change. Two means were significantly different when p < 0.05.

### 2.4 Human iPSC maintenance and cardiomyocyte differentiation

Human iPSC lines were obtained from the Wicell Research Institute (iPS IMR90-1) (https://www.wicell.org/) and the Allen Institute (https://alleninstitute.org/). Differentiation of hiPSCs to ventricular cardiomyocytes (hiPSC-CMs) employed a protocol published previously. In brief, hiPSCs were maintained on a Corning Matrigel (0.5 mg/ 6-well plate, dissolved in DMEM/F- 12 medium) cultured in mTeSR1 medium (StemCell Technologies, Vancouver, BC). Cells were passaged every 4 days using Versene solution (ThermoFisher, Waltham, MA) and were then seeded on a 6-well Matrigel-coated plate at a density of 100,000 cells cm-2 in mTeSR1 medium. The medium was changed daily, and after 3 to 4 days, when the monolayer of cells reached >90% confluency, the mTeSR1 medium was replaced with Roswell Park Memorial Institute Medium (RPMI) 1640 basal medium (ThermoFisher, Waltham, MA) plus 2% B27 without insulin supplement (ThermoFisher) containing 12 µM CHIR99021 (R&D Systems) for 24 h, followed by 5 mM IWP4 (Tocris) for 2 days without medium change. The medium was changed to RPMI 1640 plus B27 complete supplement, and then subsequently the medium was changed every 2 to 3 days. Robust spontaneous beating of the monolayers was observed by day 12.

### 2.5 Preparation and design of CRISPR-Cas9 and donor template

The single-guide RNAs (sgRNA) were prepared using online CRISPR tools including CRISPRko Tool from the Broad Institute, the CRISPR Finder from Wellcome Trust Sanger Institute, and Benchling. The sgRNA sequences were designed to target the region close to R278 residue of *TNNT2* as shown in Figure S2. They were then cloned into a pCCC vector which is based on pSpCas9(BB)-2A-GFP vector (PX458, Addgene plasmid # 48138). The pCCC vector contains the complete U6 promoter for enhanced expression in hiPSCs. The 127 bp asymmetric single-stranded donor nucleotides (ssODNs)^29, 30^ were designed (Figure S2) to include the transitions (T -> A) and (C -> T) that are seen in R278C as well as a silent mutation at the PAM site to prevent the continuous cutting and improve the efficiency of the HDR repair.

### 2.6 Transfection and Fluorescence-Activated Cell Sorting (FACS)

Human iPSCs were co-transfected with the R278C-sgRNA-pCCCplasmid and the ssODN donor template using Lipofectamine 3000 (ThermoFisher). Briefly, 2 × 105 hiPSCs were transfected with 500 ng sgRNA plasmid and 10 pmol of ssODN. Since the vector contained GFP as a reporter, the GFP+ hiPSCs were sorted as single cells using Fluorescence-Activated Cell Sorting (FACS) 48–72 hours post-transfection and were plated on Matrigel-coated plates in mTeSR122. More than 100 single-cell colonies were genotyped. To screen for on-target single mutation, genomic DNA was extracted (DNEasy, Qiagen, Mississauga, ON) and used for PCR amplification of the region around the CRISPR target sites. These amplicons were Sanger sequenced to confirm the single nucleotide mutation at the R278 *TNNT2* site using the primers described in Figure S2.

### 2.7 Protein isolation and quantification

The myocardial proteins were isolated from hiPSC-CM pellets (400k-500k cells) for both cell lines approximately 30 days after differentiation. The timely collection of the CMs was done using STEMdiff reagent from the CM dissociation kit (STEMCELL Technologies, Vancouver, BC). STEMdiff was added to each well of hiPSC-CMs, and after incubation with its neutralizing agent, the cells detached from the Geltrex matrix and were pipetted up and down for a homogenous suspension. This was followed by centrifugation of the collected cells to obtain a pellet of hiPSC-CMs. The pellet was then resuspended in lysis buffer according to the number of cells. The cells were counted using the CellDrop automated cell counter (Froggabio, ON). The lysis buffer used contained 100 mM HEPES (pH= 8-8.5) + 20% SDS, and 75 μL of were added per∼ 2 million cells. Cell lysates were then placed for 5 minutes in 95°C water bath, followed by 3 minutes on ice. To complete the lysis, the samples were sonicated twice for 30 seconds, and placed on ice between rounds. This was followed by treatment with benzonase (EMD Millipore-Sigma) at 37°C for 30 min to fragment the chromatin.

After obtaining the CM lysate, each sample was reduced with 10 mM dithiothreitol (DTT) at 37 °C for 30 min, followed by alkylation with 50 mM chloroacetamide (CAA) for 30 min in the dark. The alkylation was quenched in 50 mM DTT for 10 min at room temperature. To clean the lysates, high-resolution sample-preparation (SP3) technology was the purification approach used to isolate the proteins in the samples. This protocol is a paramagnetic bead–based approach for rapid and efficient processing of protein samples before proteomic analysis^31^. SP3 uses a hydrophilic interaction mechanism for the removal of interfering components (e.g., detergents, salts, buffers, acids, and solvents). The magnetic beads facilitate a wide range of nonselective protein binding that is enabled through the use of an ethanol-driven solvation layer on the surface of the beads. To each sample, hydrophilic and hydrophobic Sera-Mag Speed Beads (GE Life Sciences, Washington, US) were added in 1:1 ratio. Proteins were then bound to the beads with 100% ethanol (80% v/v) and washed twice with 90% ethanol.

After binding of the proteins to the beads, the samples underwent an overnight enzymatic digestion with trypsin (1:50 w/w), followed by C18 matrix midi-column clean up and elution. The BioPure midi columns (Nest Group Inc.) were conditioned with 200 μL methanol, 200 μL 0.1% Formic Acid (FA), 60% Acetonitrile (ACN), and 200 μL 0.1% Trifluoroacetic Acid (TFA). The pH of the samples was adjusted to 3 – 4 using 10% TFA prior to loading the columns. Samples were sequentially eluted 3 times with 70 μL, 70 μL, and 50 μL 0.1% FA, 60% ACN. The eluate was collected into LoBind tubes and placed into a SpeedVac to remove the organic ACN. Lastly, the samples were re-suspended in 0.1% FA (ready for MS injection). The total protein concentration for each of the purified samples was determined using Nanodrop. Mass spectrometric analyses were performed on Q Exactive HF Orbitrap mass spectrometer coupled with an Easy-nLC liquid chromatography system (Thermo Scientific). This intricate system detects the ionized peptides at the specific time that they elute out of the LC column (retention time). This time is calculated as the period between injection to detection.

In the MS sample chamber (96-well plate), the samples were completely randomized across all conditions and biological replicates. A total of 1 μg peptide per sample was injected for analysis. The peptides were separated over a three-hour gradient consisting of Buffer A (0.1% FA in 2% acetonitrile) and 2%-80% Buffer B (0.1% FA in 95% acetonitrile) at a set flow rate of 300 nL/min. The computerized MS setup produces a pattern spectrum of masses based on the abundance and structure of the detected ionized molecules.

The raw mass-to-charge (M/Z) data acquired from the Q Exactive HF were searched with MaxQuant (MQ)-version 2.0.0.0 (Max Planck Institute, Germany), and Proteome discoverer software (Thermo Fisher Scientific, CA, USA), using the built-in search engine, and embedded standard DDA settings. The false discovery rate for protein and peptide searches was set at 1%. Digestion setting were set to trypsin. Oxidation (M) and Acetyl (N-term) were set as dynamic modifications. Carbamidomethyl (C) was set as fixed modification, and Phosphorylation (STY) was set as a variable modification. For the main comparative protein search, the human proteome database (FASTA) was downloaded from Uniprot (2022_06; 20,365 sequences) and used as the reference file. Common contaminants were embedded from MaxQuant. Peptide sequences that were labeled as “Potential contaminant/REV” were excluded from the final analysis.

### 2.8 Optical mapping for action potential and calcium transient recordings

Prior to imaging, monolayers of hiPSC-CMs ± R278C^+/-^ were perfused with IMDM supplemented with a physiological solution (final concentrations in mM): 140 NaCl, 3.6 KCl, 1.2 CaCl_2_, 1 MgCl_2_, 10 HEPES and 5.5 D-glucose for optical mapping as described^20, 27, 32, 33^. In brief, the cells were incubated with 15 μM of the potentiometric dye RH-237 (Molecular Probes, Eugene, OR) for 45 minutes. After incubating the hiPSC-CMs with RH-237, 5 μM of the Ca^2+^-sensitive dye Rhod-2AM (Molecular Probes), 20 μM blebbistatin (a myosin ATPase inhibitor (Sigma-Aldrich)) were added to the culture dish. Blebbistatin inhibits contraction and reduces motion artifact with no detectable effect on the electrophysiology properties^34, 35^. For imaging, the hiPSC-CMs were excited by 532 nm LEDs. RH-237 and Rhod-2 emissions were monitored using a >710 nm long-pass and 565-600 nm band-pass filters, respectively, at 100 fps at 37°C. The spectral properties allow both dyes to be fluoresced using 532 nm and their respective emissions to be separated by dichroic mirrors. Both signals were captured with a single Hamamatsu ORCA Flash 4 digital CMOS camera by incorporating an optical image splitter. Electrical field stimulation was applied using stainless steel electrodes placed ∼1 cm apart in the imaging chamber. The hiPSC-CMs harboring the *TNNT2* heterozygous mutation R278C^+/-^, and WT were stimulated for 20 seconds each at four stimulation frequencies (55, 65, 75, and 100 beats/min). Data were analyzed using custom-built software in IDL (Exelis Visual Information Solutions, McLean, VA)^27, 32, 33^.

### 2.9 Contractile parameters assessment using SarcTrack

The contractile properties of both hiPSC-CMs ± R278C^+/-^ were studied using SarcTrack^22^ about 30 days after differentiation. SarcTrack facilitates high throughput contractile measurements of 2D monolayers of hiPSC-CMs for disease modeling and large-scale pharmaceutical drug screening. This MATLAB algorithm fits each sarcomere to a double wavelet with 6 definable parameters including the distance between wavelet peaks as a function of time, constraint length of each wavelet, width of each individual wavelet, number of angles that the wavelet test used to find the best fit to the z-disc, distance between grid points to avoid refitting same sarcomere, and the neighborhood around each grid where the best fit for the wavelet pair can be determined. SarcTrack was used to measure the real time contractility of thousands of sarcomeres by running multiple videos in parallel. The images were acquired with a Hamamatsu Orca Flash scMOS camera at 100 frames per second (fps) to measure contractility in hiPSC-CM monolayers cultured on PDMS matrices of 5 kPa tensile strength and stimulated at 1 Hz at 37℃. To image fluorescently labeled sarcomeres, the α-actinin 2-GFP tagged hiPSC-CMs (Allen cell line (AICS-0075)) were used. The images acquired at 100 fps resulted in good signal-to-noise ratio based on a root mean square (RMS) analysis.

On day 14 of differentiation, WT and R278C^+/-^ variant hiPSC-CMs were replated on a Polydimethylsiloxane (PDMS) substrate with a stiffness of 5 kPa to provide a physiological environment for the cells and to increase their maturation. To create a PDMS substrate with 5 kPa elastic modulus, commercially available PDMS, Sylgard 527 gel (equal weights of part A and part B) was used. The surface of PDMS substrate was coated with fibronectin (50 µg/ml) and hiPSC-CMs were seeded with the concentration of 200,000 cell cm^-2^. Two weeks after replating, the samples were stimulated at 1 Hz frequency and 5 second videos were captured at 100 fps with an ROI of 1024*1024 pixels. A Nikon Ti2 microscope was used with 100x oil immersion objective and several videos were captured from different parts of the substrate. Next, by defining the appropriate SarcTrack parameters, multiple videos were run simultaneously. To run more than 100 videos at the same time, the files were uploaded to the computational cluster at the Digital Research Alliance of Canada. This advanced research computing system allows one to specify different numbers of CPUs and decreases the computational cost. In this study, 32 CPUs were defined to run more than 100 videos in about 10 hours. The measured contractility parameters included the percentage of sarcomere shortening, time to half contraction, time to half relaxation, maximum velocity of contraction, and maximum velocity of relaxation of WT and R278C^+/-^ variant hiPSC-CMs. To study the effect of mavacamten on the contractility parameters, R278C^+/-^ variant hiPSC-CMs were treated with 0.3, 1, and 3 µM mavacamten for 15 min before capturing videos.

### 2.10 Statistical analysis

Average data are provided as mean ± standard error of mean (SEM). The generated MS raw data were filtered and transformed to log2 scale. This was followed by quantile normalization for all samples. The proteomic profile for the HCM variant was subject to comparative analysis against the isogenic control cell line. The protein groups file obtained from the DDA MQ search was used for all the quantitative statistical analysis based on the detected intensities. All plots, visualizations and statistical comparisons were done using R and Perseus (version 1.6.15.0) software based on unpaired two tailed Student’s t-test of significance. The level of significance (p-value < 0.05) and number of data points (n) are indicated in each figure.

## 3. RESULTS

### 3.1 Computational potential energy minimization of WT and R278C^+/-^ cardiac thin filament

Potential energy minimizations of our Rosetta models were carried out for the cardiac thin filament with and without and the R278C cTnT variant (Figure 1). The R278C^+/-^ potential energy minimization analysis suggests that the substitution of a positively charged arginine to a more hydrophobic cysteine may cause more favorable local interactions. The residues in the actin and tropomyosin molecule that are within proximity of residue 278 of cTnT create a slightly positively charged environment. Therefore, the native R278, which is positively charged, may be less favorable to exist in this environment due to the charge repulsion compared to a mutated R278C. If this repulsion of an R278 residue is necessary for normal function (i.e., to provide some flexibility in the local region), we propose that the R278C mutation may disrupt this function due to the lack of the repulsion. In addition, there was no nearby cysteine from actin or Tm that could form any potential disulfide bridges with R278C.

With the cutoff set at 10 Å, five residues of Tm were within the range of the CG or the SG atom for all eight models: Arg167, Val170, Ile171, Ser174 and Arg178 (chain P/W). At this cutoff, the four 6KN7 CG/SG atoms of the models are within range of residues within the closest actin (F/G), whereas the four 6KN8 CG/SG atoms are not within range of any actin molecules. Analysis of this shows that the mutated R278C^+/-^ models of 6KN7 are closer, by about 2 Å, to Arg147 and Thr148 of the actin molecule. Two representations of residue 278 (one in the wildtype actin chain F the second in the variant form) in its environment (with proximal actin/tropomyosin residues labeled) are shown in Figure 1)^36, 37^.

With the cutoff set at 10 Å, five residues of tropomyosin were within the range of the CG or the SG atom for all eight models: Arg167, Val170, Ile171, Ser174 and Arg178 (chain P/W). At this cutoff, the four 6KN7 CG/SG atoms of the models are within range of residues within the closest actin (F/G), whereas the four 6KN8 CG/SG atoms are not within range of any actin molecules. Analysis of this shows that the mutated R278C^+/-^ models of 6KN7 are closer, by about 2 Å, to Arg147 and Thr148 of the actin molecule. Two representations of residue 278 (one in the wildtype actin chain F the second in the variant form) in its environment (with proximal actin/tropomyosin residues labeled) are shown in Figure 1^36, 37^.

### 3.2 Ca^2+^ sensitivity and dissociation rates in human cardiac reconstituted thin filaments (hcRTF)

lterations to myofilament Ca^2+^ sensitivity are a hallmark of HCM-associated variants and can explain pathological changes seen in HCM such as hypercontractility and metabolic dysfunction^38^. To study this, steady-state fluorescence measurements were carried out on WT and R278C^+/-^ hcRTFs, and Ca^2+^-dependent IAANS fluorescence increase was observed for all hcRTF constructs. The R278C^+/-^ variant showed a statistically significant increase of thin filament Ca^2+^ sensitivity by ∼2.3 fold (p<0.05) (Figure 2). Stopped-flow fluorescence spectroscopy was used to determine the Ca^2+^ dissociation rates from the hcRTF constructs. Upon the addition of EGTA, the Ca^2+^ rapidly dissociated from cTnC, inducing a cTnC conformational change in which the fluorescence change was recorded. As Ca^2+^ was stripped from cTnC by EGTA, a decrease in IAANS fluorescence was observed for all hcRTF constructs. hcRTF harboring R278C^+/-^ show a significant decrease of k_off_ compared to WT, significantly increasing the Ca^2+^ sensitivity by ∼1.4 fold (p<0.05) (Figure 2). This explains, at least in part, the increase of Ca^2+^ sensitivity conferred by the hcRTF harboring the R278C^+/-^ variant in the steady-state experiments.

**Figure 2.**
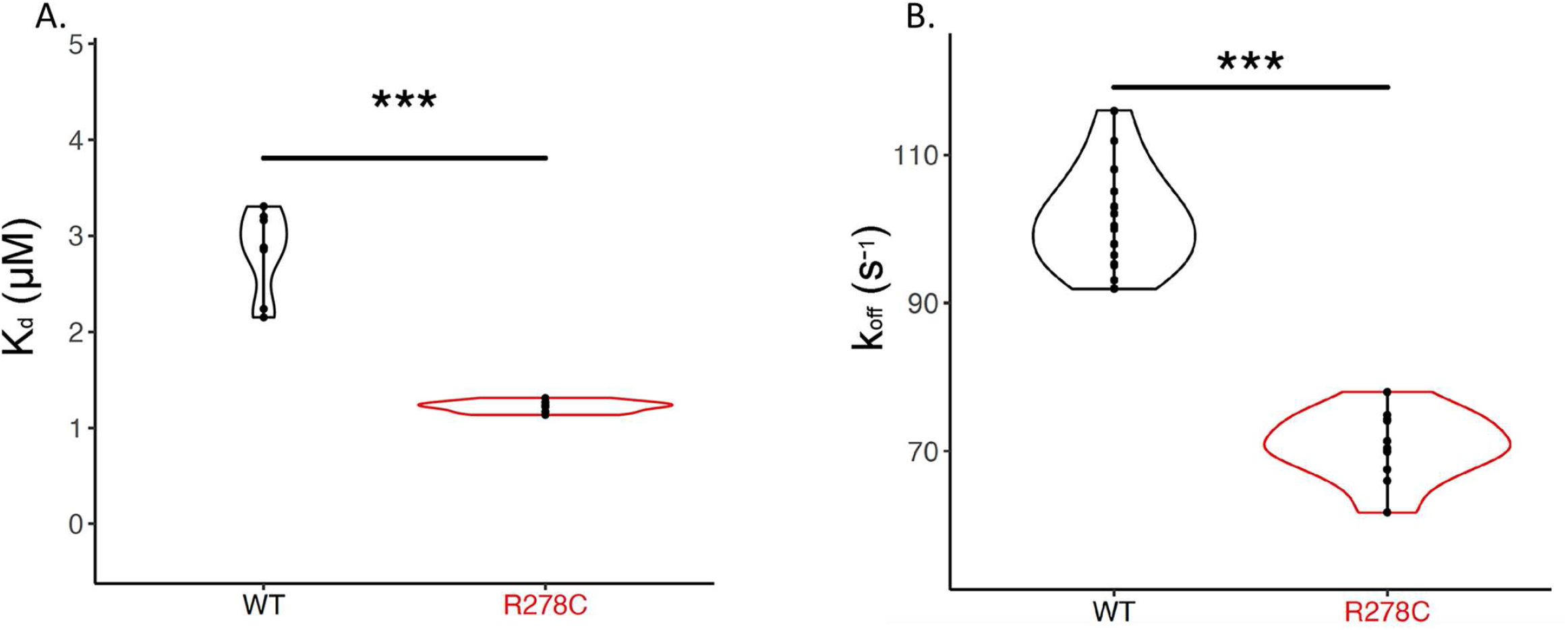
Steady-state and stopped-flow analysis of Ca^2+^ kinetics in human cardiac reconstituted thin filaments (hcRTF). Panel A. The Ca^2+^ sensitivity (expressed in K_d_ (µM)) of the WT and R278C^+/-^ hcRTF was determined by steady-state fluorescence spectrometry. The R278C significantly decreased the K_d_ compared to the WT counterpart by ∼2-fold, indicating that this variant increases the Ca^2+^ sensitivity significantly in the hcRTF biochemical system. Panel B. A stopped- flow instrument was used to determine the Ca^2+^ dissociation rates (expressed in k_off_ (s^-1^)) from the hcRTF. Upon the addition of EGTA, the Ca^2+^ was rapidly removed from TnC, inducing a conformational change in cTnC within hcRTF in which the fluorescence change was recorded. The R278C^+/-^ hcRTF exhibited a decrease in the k_off_ by ∼1.5 fold, compared to the WT counterpart, suggesting that the increase in Ca^2+^ sensitivity observed in the steady state is largely due to a slower Ca^2+^ off-loading rate. *p<0.05, **p<0.05, ***p<0.005.

### 3.3 Protein quantification and post-translational modification characterization

Previous studies have shown that HCM can largely alter the proteomic profile of the heart, highlighting the extensive underlying pathological remodeling^39, 40^. To investigate the proteomic alterations caused by R278C, we obtained the full proteomic profile of *TNNT2* R278C^+/-^ and isogenic control hiPSC-CMs using shotgun, bottom-up mass spectrometry. Here, we found 102 of 2327 proteins to be statistically (1% FDR) more abundant in WT and 198 proteins to be more abundant in R278C^+/-^ cells (Figure 3A). Next, we examined individual proteins of interest involved in control of contractility, structural integrity, metabolism, and crucial cardiac remodeling pathways (Figure 3B-D). The variable profile was further probed among individual protein groups. Even though no significant difference was noted in most sarcomeric proteins, the myosin heavy chain showed a dissimilarity between *TNNT2* R278C^+/-^ and isogenic control hiPSC-CMs. As seen in Figure 3B, the dominant form of myosin heavy chains (MHC), Myosin Heavy chain 7 (MYH7; βMHC) with relatively slow ATPase activity, did not exhibit any quantitative changes between the two groups. However, the other key paralog, MYH6 (αMHC) with relatively fast ATPase activity, was significantly over-expressed in the *TNNT2* R278C^+/-^ hiPSC-CMs. Also, the myosin light chains (MYLs), particularly MYL2, showed a significant decrease in the R278C-harboring CMs relative to the wildtype cells (Figure 3B).

**Figure 3.**
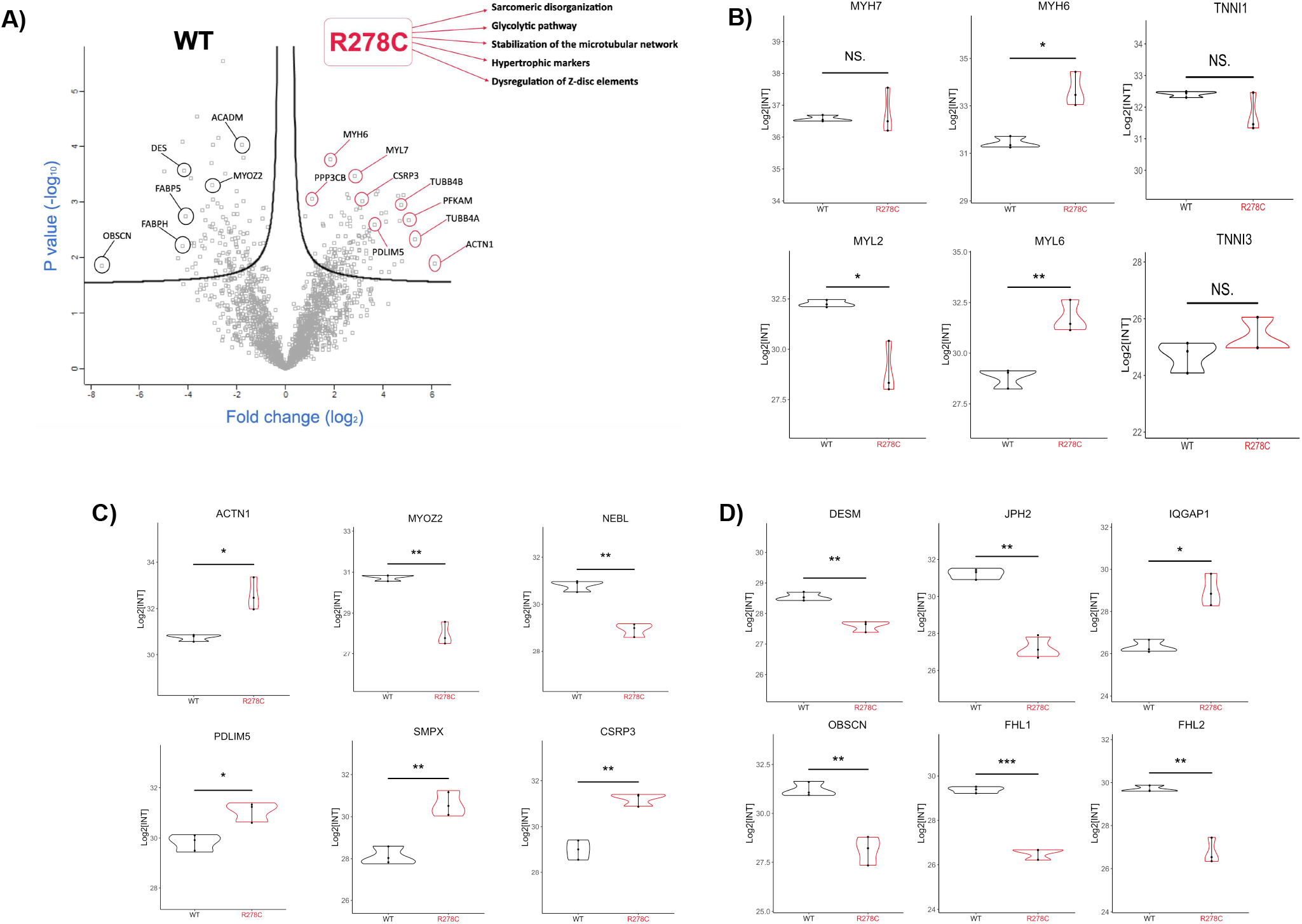
MS proteomic abundance profiles. Panel A. Volcano plot comparison of WT vs. R278C^+/-^ on the protein expression levels. The right side of the plot (red) reflects upregulated proteins in R278C^+/-^ hiPSC-CMs, and the left side (black) shows the proteins that were detected in higher quantity in the WT hiPSC-CMs. The FDR threshold was set to 0.01, and any data points beyond that were considered as statistically significant. Gene ontology (GO) annotation terms summarize the top cross search results in reference to KEGG, Elsevier, Jensen, and GO biological processes databases. Panels B-D. Violin plot representations of individual proteins that exhibited significant changes between the R278C^+/-^ vs. WT and separated according to their functional groups. Panel B. Thick filament proteins: MYH7 (myosin heavy chain-7), MYH6 (myosin heavy chain-6), MYL2 (myosin regulatory light chain ventricular isoform), and MYL6 (myosin regulatory light chain-6). Panel C. Z-disc proteins: ACTN1 (Alpha actinin-1), MYOZ2 (Myozenin-2), NEBL (Nebulette), PDLIM5 (PDZ and LIM domain protein-5), SMPX (Small muscular protein), and CSRP3 (Cysteine and glycine-rich protein-3). Panel D. Cardiac remodelling proteins: DESM (Desmin), JPH2 (Junctophilin-2), IQGAP1 (Ras GTPase-activating-like protein), OBSCN (Obscurin), FHL1 (Four and a half LIM domains protein-1), and FHL2 (Four and a half LIM domains protein-2). *p<0.05, **p<0.05, ***p<0.005; two-tailed unpaired t-test.

Proteins important for the structural integrity and organization of the Z-disc showed variable translational profiles between the R278C^+/-^ and WT CMs (Figure 3C). α-actinin, a major protein involved in the crosslinking of actin filaments that is disrupted in HCM^41^, was significantly increased in *TNNT2* R278C^+/-^ hiPSC-CMs. Meanwhile, other protein groups responsible for the structural integrity and stability of the Z-disc, such as MYOZ2 (Myozenin2), NEBL (Nebulette), and SMPX (small muscle protein)^42, 43^, were significantly downregulated in R278C^+/-^ hiPSC-CMs, which indicates a further exacerbation of the Z-disc inflexibility and disorganization. Microtubule related proteins such as tubulin β-4B chain (TUBB4B), which play a crucial role in signal transduction and contractility^44^, were significantly increased in R278C^+/-^ hiPSC-CMs. Other structural proteins responsible for the overall integrity of the cardiomyocyte also showed significantly lower protein expression in R278C^+/-^ hiPSC-CMs (Figure 3C and D). These include NEBL (Nebulette), which functionally links sarcomeric actin to the desmin intermediate filaments in the sarcomeres^45^, JPH2 (Junctophilin2), which provides a structural bridge between the plasma membrane and the sarcoplasmic reticulum and is required for normal excitation-contraction coupling in cardiomyocytes ^46^, OBSCN (Obscurin), which is involved in the assembly of myosin into sarcomeric A bands ^47^, FHL 1&2 (Four and a half LIM domains), which negatively regulate the calcineurin/NFAT signaling pathway in cardiomyocytes ^38^, and DES (Desmin), a key structural intermediate filament required for maintaining normal structure of sarcomeres and interconnecting the Z-discs^39^. These data suggest significant changes in structural integrity and Z-disc disorganization which may underlie the myofibrillar disarray seen in HCM.

Lastly, a metabolic shift in the proteomic profile was observed in R278C^+/-^ hiPSC-CMs, in which glycolytic proteins were significantly up-regulated, including GLUT1 (insulin-independent glucose transporter protein type 1), PFKAM (ATP-dependent 6-phosphofructokindase muscle type), and PGAM1 (phosphoglycerate mutase 1) (Figure 4A). Meanwhile, major fatty-acid metabolism and β-oxidation proteins, such as ACADS, ACADM, and FABPH, were significantly downregulated in R278C^+/-^ hiPSC-CMs (Figure 4B). These data suggest that R278C may cause a switch to a fetal-like metabolic profile, which has been previously shown in HCM^48^. Additionally, a major metabolic hallmark in HCM is impaired energetics and inefficient ATP utilization^49^. We found CKM (creatine kinase, muscle), CKMT2 (mitochondrial creatine kinase), and ATPD (ATP Synthase F1 Subunit Delta) to be significantly decreased in R278C^+/-^ hiPSC-CMs (Figure 4C), suggesting disrupted cardiac bioenergetics through impaired ATP production and utilization.

**Figure 4.**
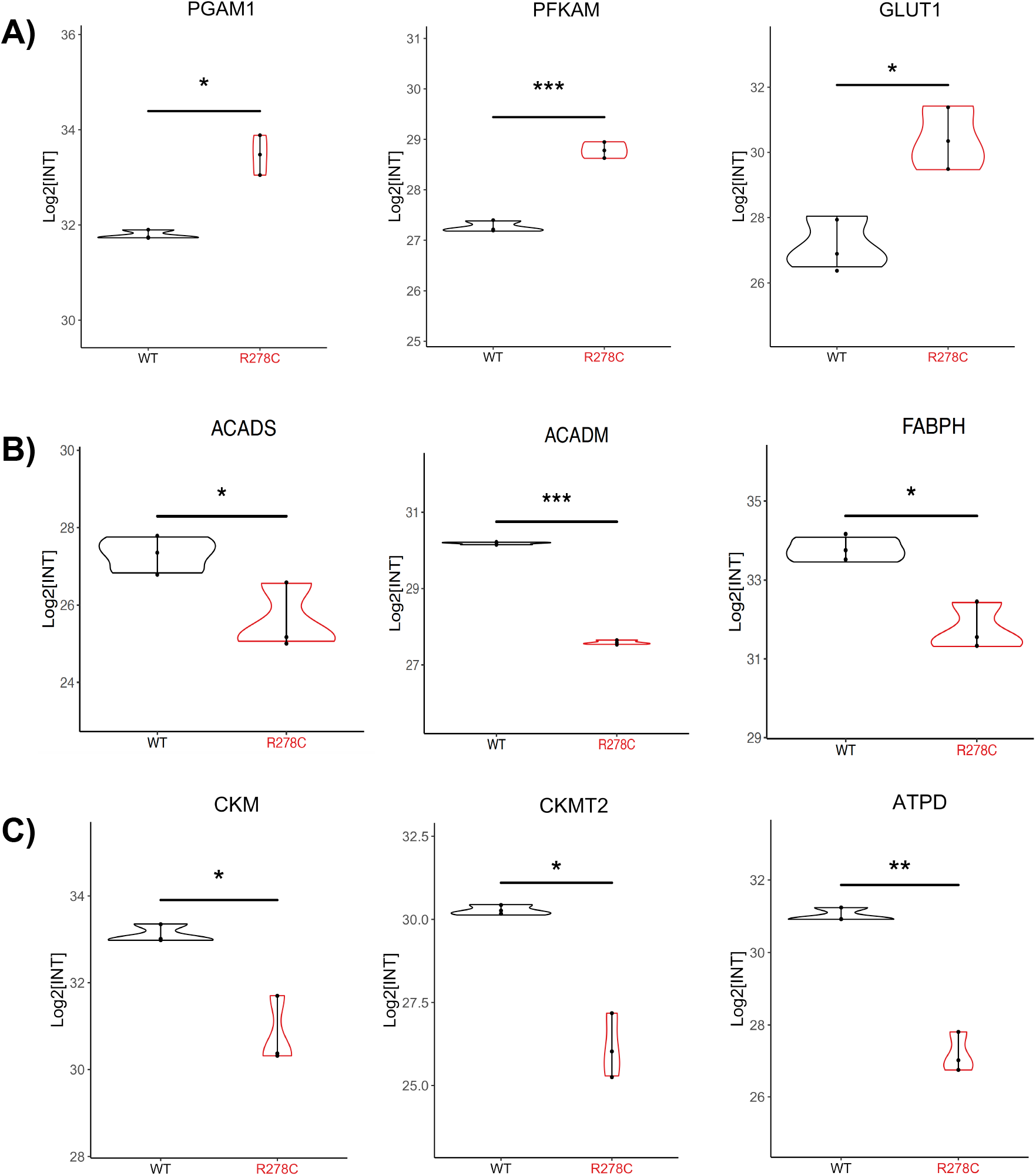
MS proteomic abundance profile of metabolic and energy production proteins. Panel A. PGAM1 (Phosphoglycerate mutase-1), PFKAM (ATP-dependent 6-phosphofructokinase, muscle type), GLUT1 (Solute carrier family 2, facilitated glucose transporter member-1). Panel B. ACADM (Medium-chain specific acyl-CoA dehydrogenase), ACADS (Short-chain specific acyl-CoA dehydrogenase), FABPH (Fatty acid binding protein, heart). Panel C. CKM (Creatine kinase M-type), CKMT2 (Creatine kinase mitochondrial type 2), and ATPD (ATP synthase subunit delta, mitochondrial). (* p-value< 0.05, ** p-value <0.005, ***p-value<0.0005; two-tailed unpaired t-test.)

In addition to the differences in protein expression, significant alterations were observed on the post translational level, particularly phosphorylation (Figure 5). We found significant dephosphorylation of NFATC2 in *TNNT2* R278C^+/-^ hiPSC-CMs. As the NFAT-Calcineurin pathway is a major contributor to pathological hypertrophy and remodeling^50^, this mechanistically links R278C to its potential downstream effects. Moreover, we found MYBPC3 and MYL2, both of which are important regulators of the super relaxed (SRX) and disordered relaxed (DRX) states of the myosin heads^51^, to be differentially phosphorylated at sites Ser284 and Ser15, respectively, with higher phosphorylation in R278C^+/-^ hiPSC-CMs. These data suggest a favored transition to the DRX state with R278C, which is a pathological transition seen in HCM.

**Figure 5.**
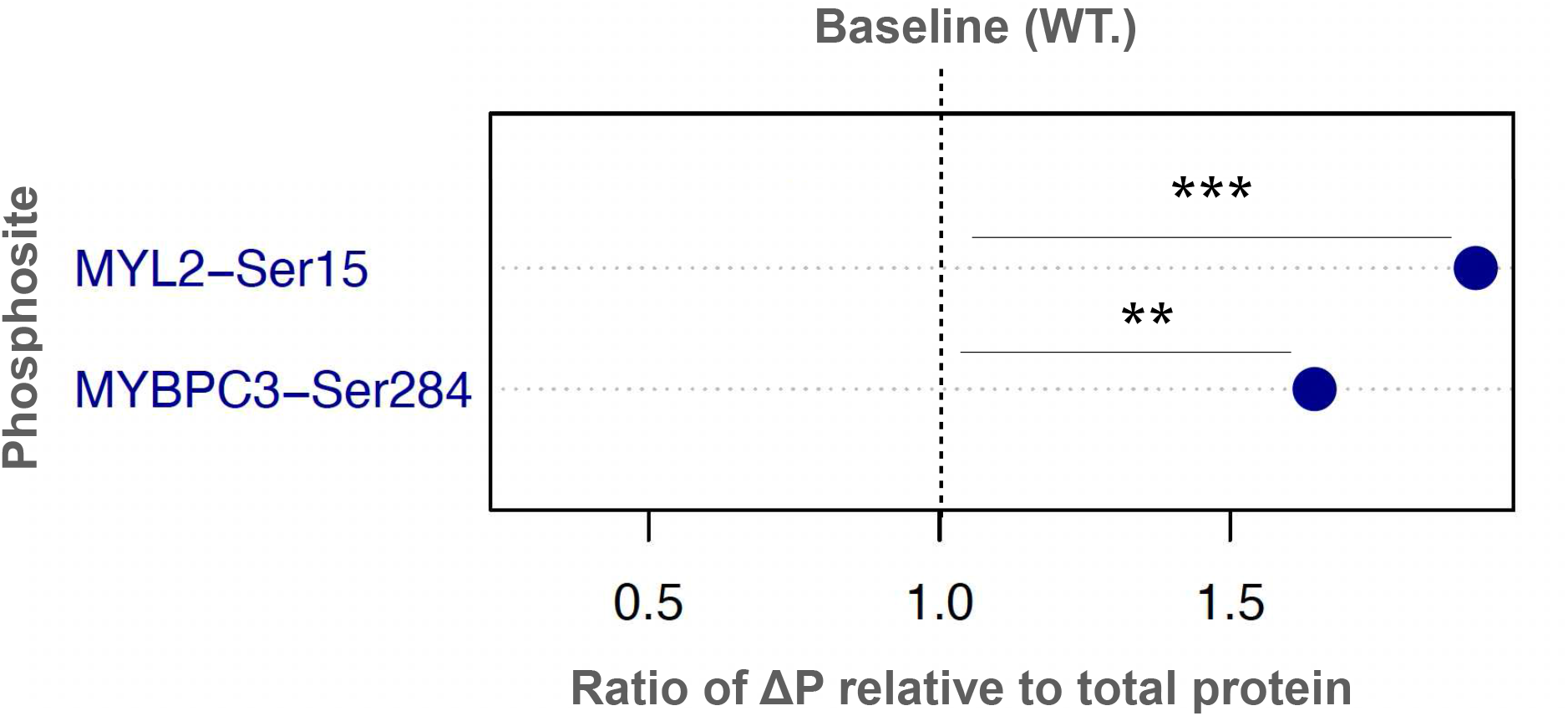
Phosphorylation profile of six selected phosphosites in R278C^+/-^ vs. WT hiPSC-CMs. Dot plot representation of the phosphorylation change relative to the total protein expression. The degree of WT phosphorylation was used as the baseline to compare the phosphorylation trends in the chosen phosphosites in the R278C^+/-^ hiPSC-CMs.

### 3.4 Electrophysiological studies in hiPSC-CM monolayers

Using optical mapping on 2D monolayers of R278C^+/-^ and isogenic control hiPSC-CMs, we investigated the effect of various stimulation frequencies on the membrane voltage (Vm) and cytosolic Ca^2+^ transients. Vm and Ca^2+^ transients were determined from the fluorescence emitted by RH-237 and Rhod-2, respectively. Figure 6A shows representative traces of Vm and Ca^2+^ transients from monolayers of hiPSC-CMs electrically stimulated at 55, 65, and 100 beats per minute (bpm). The action potential and Ca^2+^ transients for both control (A1, A2) and R278C^+/-^ (A4, A5) hiPSC-CMs show that both groups entrained normally to 55 and 65 bpm. However, while the control hiPSC-CMs showed normal entrainment to 100 bpm (A3), R278C^+/-^ hiPSC-CMs developed alternans-like irregular and arrhythmogenic voltage and Ca^2+^ transients at this frequency (A6). None of the control hiPSC-CM monolayers exhibited abnormal V_m_ or Ca^2+^ transients at 100 bpm. This suggests that fast stimulating rates can be arrhythmogenic in R278C^+/-^, which may underlie the higher risk for ventricular arrythmias and sudden death.

**Figure 6.**
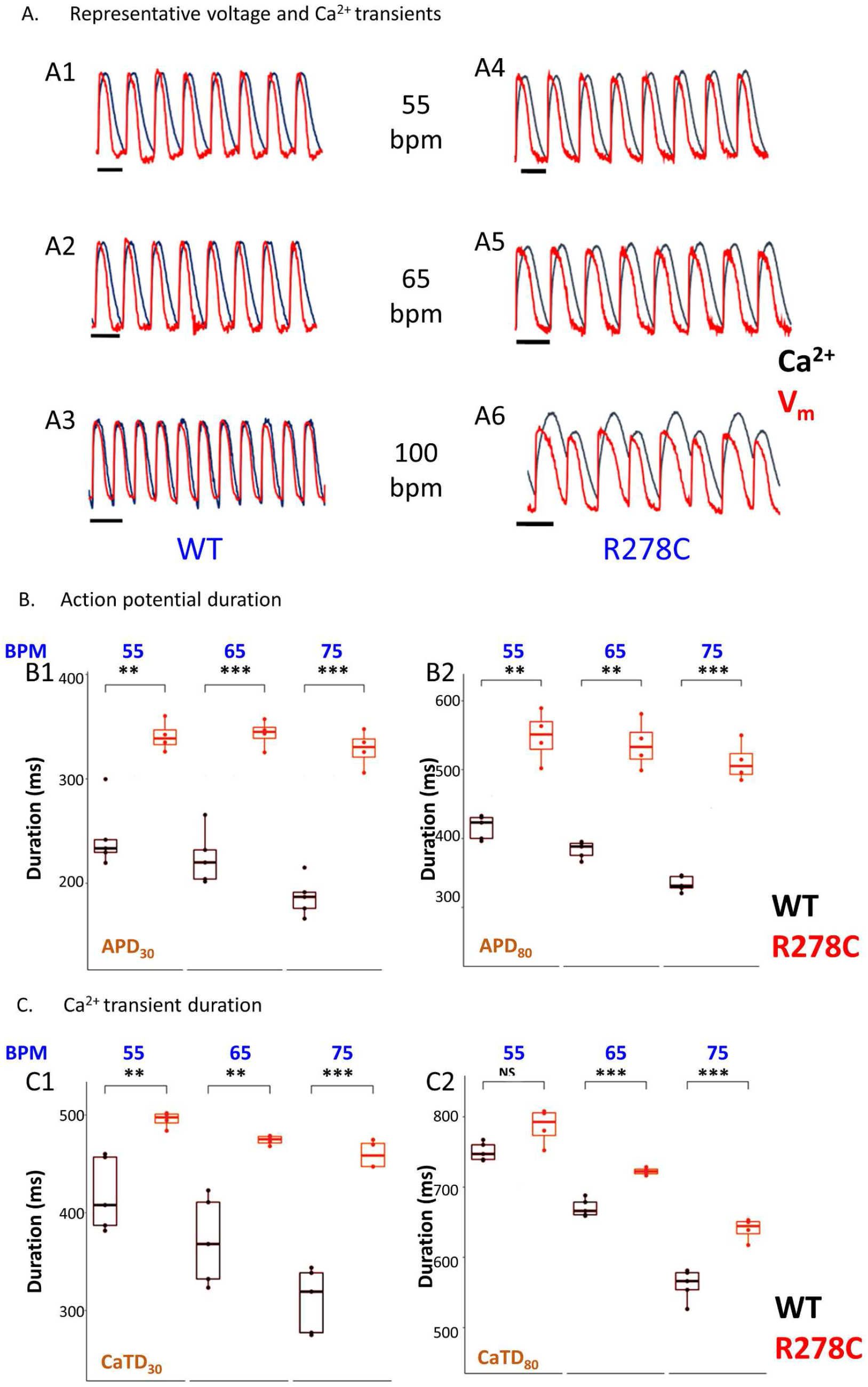
Optical mapping (OM) of membrane voltage (Vm) and calcium (Ca^2+^) transients in 2D monolayers of hiPSC-CMs. Panel A. Representative OM traces of V_m_ (red) and Ca^2+^ (black) transients in WT (A1 to A3) and R278C^+/-^ (A4 to A6) hiPSC-CMs at three different stimulation frequencies 55, 65, 100 bpm in the top, middle, and bottom traces, respectively. WT hiPSCs responded normally to all stimulation frequencies, however, about 50% of R278C^+/-^ hiPSC-CMs exhibited voltage and Ca^2+^ alternans at frequencies above 75 bpm. Panel B. Statistical analysis of action potential duration of WT (black) and R278C^+/-^ (red) hiPSC-CMs as a function of stimulation frequencies over the range of 55 to 75 BPM. Panels B1 and B2 show that both the APD_30_ and APD_80_ were significantly longer in R278C^+/-^ vs. WT at all stimulation frequencies. Panel C1 shows that the CaTD_30_ was significantly longer in R278C^+/-^ vs. WT at all stimulation frequencies. Panel C2 exhibits the longer CaTD_80_ in R278C^+/-^ vs WT hiPSCs at 65 and 75 bpm. (* p-value< 0.05, ** p-value <0.005, ***p-value<0.0005)

We also examined action potential durations at 30% (APD_30_; B1) and 80% (APD_80_; B2) of repolarization as a function of stimulation frequency (55, 65, and 75 bpm) in both control and *TNNT2* R278C^+/-^ hiPSC-CMs (Figure 6B). Similarly, we examined Ca^2+^ transient duration at 30% (CaTD_30_; C1) and 80% (CaTD_80_; C2) of repolarization at the same stimulation frequencies (Figure 6C). These frequencies were chosen as 55 bpm was higher than the intrinsic beating rate and 75 bpm was below frequencies that induced any observable form of arrhythmia. Both APD_30_ and APD_80_ were significantly longer in R278C^+/-^ hiPSC-CMs relative to controls at all frequencies (Figure 6B1 and B2). CaTD_30_ was significantly prolonged in R278C^+/-^ hiPSC-CMs compared to WT hiPSC-CM at all stimulation frequencies (Figure 6C1), while CaTD_80_ was significantly prolonged at 65 and 75 bpm only (Figure 6C2). Moreover, we found that APD and CaTD significantly shortened with an increase in stimulation frequencies in control hiPSC-CMs, consistent with the concept of restitution. Contrarily, R278C^+/-^ hiPSC-CMs did not exhibit any indication of APD shortening under these experimental conditions, but did, however, display significantly (p<0.05) shorter Ca^2+^ transient durations as reflected by reduced CaTD_30_ and CaTD_80_ at 75 vs. 55 bpm. Collectively, R278C^+/-^ disrupts APD and Ca^2+^ transient dynamics, creating electrical instability at higher frequencies.

### 3.5 R278C^+/-^ demonstrates hypercontractility which is alleviated by mavacamten

Hypercontractility and abnormal sarcomeric shortening are classical features in HCM that drive pathogenesis. To examine the contractile properties of R278C, we employed the newly developed Matlab-based algorithm, SarcTrack, to quantify sarcomeric shortening in α-actinin 2-GFP tagged hiPSC-CMs. The goodness of double wavelet fitting to the sarcomere is shown in Figure 7A. Contractile parameters were measured while the monolayer of hiPSC-CMs was paced at 1 Hz. The percentage of maximal sarcomere shortening was significantly higher in R278C^+/-^ (14.1 ± 0.5%) compared to control (12.8 ± 0.9%) hiPSC-CMs (Figure 7C1). Dramatic increases in the time-to-half maximal contraction and relaxation were also observed in R278C^+/-^ compared to control hiPSC-CMs (Figures 7B1 and C2), which reflected the hypercontractility of R278C^+/-^ variant. However, no significant changes were observed in either the maximum velocity of the relaxation or contraction (Figure 7B2 and C3). We next investigated the effect mavacamten on contraction and relaxation parameters in R278C^+/-^ hiPSC-CMs using various concentrations (0.3, 1, and 3 µM). We found that 1 and 3 µM mavacamten caused a significant decrease in the maximum velocity of contraction and the maximum percentage of sarcomere shortening (Figure 8A), and a significant decrease in the maximum velocity of relaxation R278C^+/-^ hiPSC-CMs (Figure 8B). Thus, mavacamten may be a promising therapeutic for HCM patients carrying variants in thin filament proteins such as *TNNT2* R278C^+/-^.

**Figure 7.**
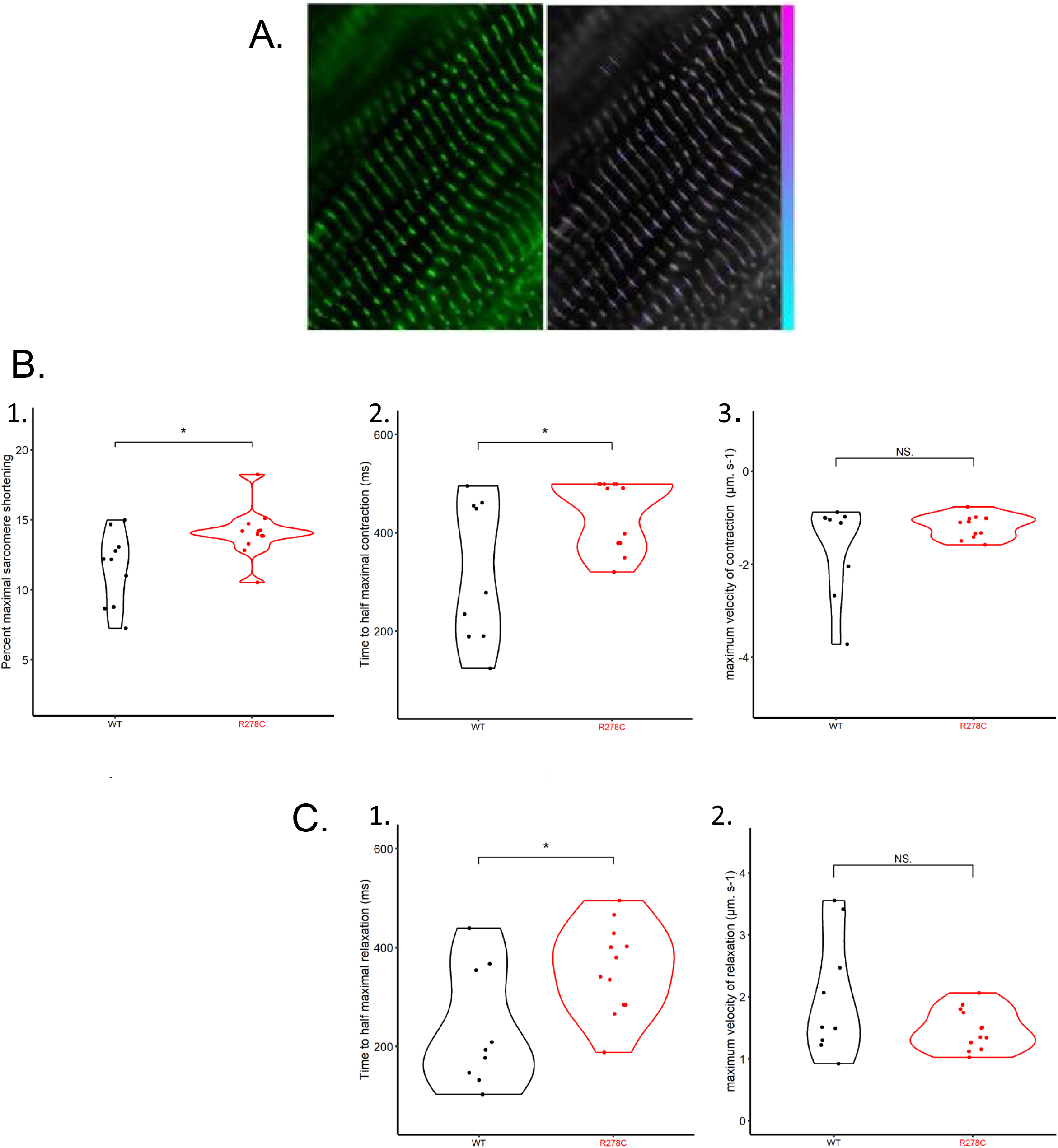
SarcTrack analysis of WT and R278C^+/-^ hiPSC-CM contractile properties. Panel A. GFP-tagged α-actinin hiPSC-CMs cultured on 5 kPa PDMS (left) and SarcTrack wavelet fitting (right). Panel B. The relaxation of WT (black, n=9) and R278C^+/-^ (red, n=12) hiPSC-CMs cultured on 5 kPa PDMS, with respect to: B1) time-to-half maximal relaxation (ms). B2) the maximum velocity of relaxation (µm.s^-1^). Panel C. The contractile parameters of WT (black, n=9) and R278C^+/-^ (red, n=12) hiPSC-CMs cultured on 5 kPa PDMS and stimulated at 1 Hz with respect to: C1) percent maximal shortening; C2) time to half maximal contraction; C3) maximum velocity of contraction (µm s^-1^). (* p-value< 0.05, ** p-value <0.005, ***p-value<0.0005)

**Figure 8.**
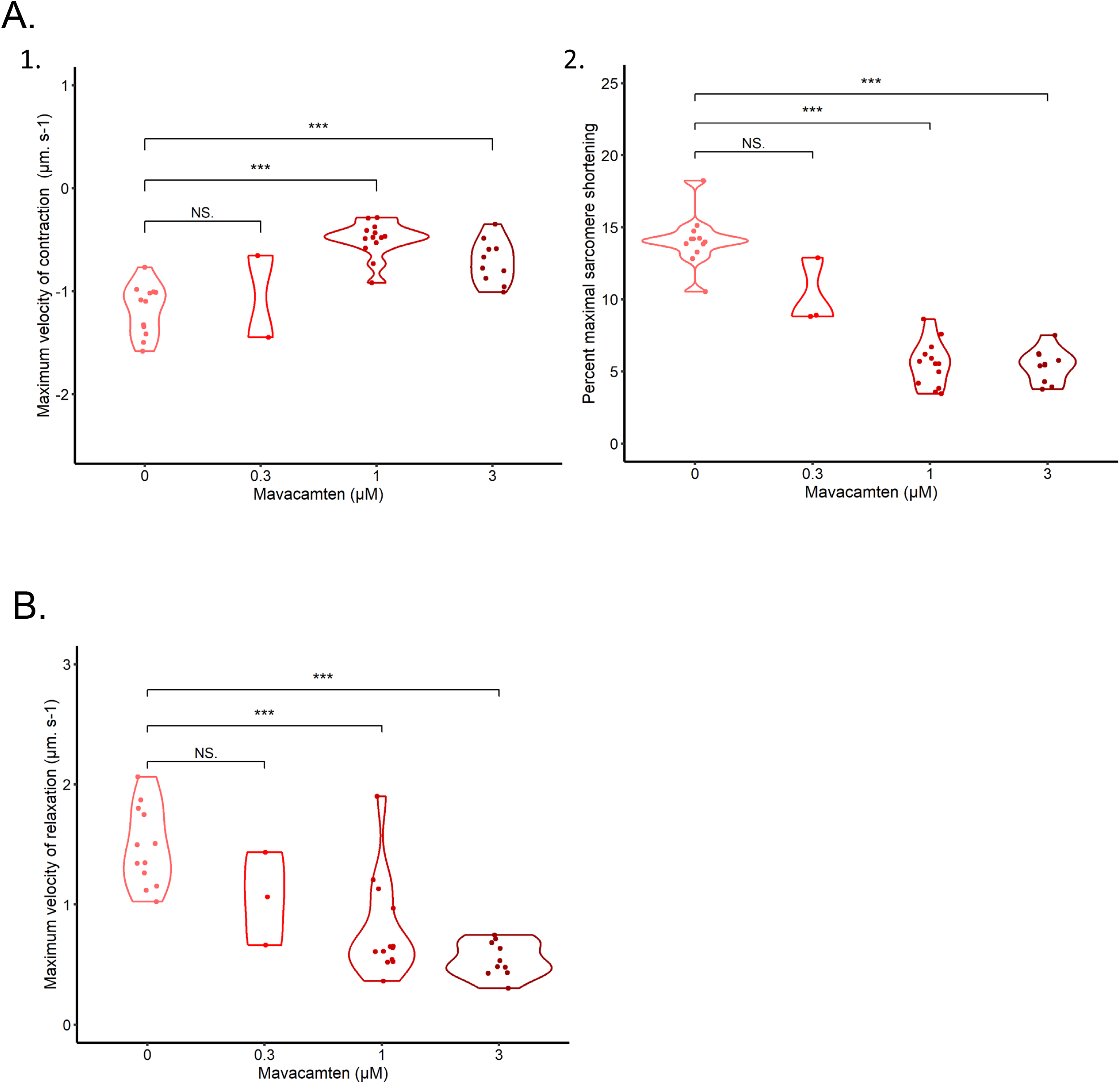
SarcTrack analysis of the effect of mavacamten on R278C^+/-^ hiPSC-CM contractile properties. Panel A. The effect of mavacamten on the contractile parameters of R278C^+/-^ (red, n=37) hiPSC-CMs cultured on 5 kPa PDMS and stimulated at 1 Hz with respect to: A1) maximum velocity of contraction (µm. s^-1^) and A2) percent maximal shortening. Panel B. The effect of mavacamten on the relaxation of R278C^+/-^ (red, n=37) hiPSC-CMs cultured on 5 kPa PDMS, with respect to the maximum velocity of relaxation (µm. s-1). (* p-value< 0.05, ** p-value <0.005, ***p-value<0.0005)

## 4. DISCUSSION

In this study, we generated in *TNNT2* R278C^+/−^ hiPSC-CMs using CRISPR/Cas9 and investigated the pathological consequences of this variant on the proteome and function of hiPSC-CMs Hypercontractility was observed in R278C^+/−^ hiPSC-CMs and mavacamten was found to alleviate it. Furthermore, an increase in Ca^2+^ sensitivity was observed in the R278C^+/-^ hcRTF model. Alterations of AP dynamics and Ca^2+^ handling in R278C^+/−^ CMs were shown to be proarrhythmic at higher stimulation frequencies. Remarkably, the proteome and contractility were dramatically impacted by the insertion of a point mutation in one allele of troponin T and likely has profound consequences in the disease phenotype. Our findings are indicative of potential pathological remodeling in R278C^+/−^ occurring in the early developmental stages of life.

### 4.1 R278C dynamics and Ca^2+^ sensitivity

Due to the highly flexible nature of cTnT, the molecular structure of most regions of cTnT remain unresolved except for residues 199-272 that are included in the high-resolution troponin core domain crystal structure (PDB: 4Y99)^36^ and recently determined thin filament cryo-EM structures (PDB: 6KN7, PDB: 6KN8). The potential energy minimization data for R278C^+/-^ suggests that the cysteine may experience slightly less charge repulsion with nearby residues from actin and Tm. As shown in Figure 1, these nearby residues in actin and tropomyosin create a positively charged microenvironment. We speculate that the charge repulsion between the positively charged native R278 and the local environment might be necessary for its normal function, in which the inherent dynamics and flexibility is required for the molecular movement that is propagated along the thin filament in response to the Ca^2+^ binding to and dissociation from the troponin complex. Previous in vitro functional analyses show that R278C alters Ca^2+^ sensitivity and actin myosin cross-bridging in rabbit, porcine, and human cardiac preparations ^17, 52–54^. Consistently, our hcRTF models show a similar magnitude of Ca^2+^ sensitization, corroborating the impact of R278C on myofilament dynamics. This is in contrast to studies in R278C^+/-^ transgenic mice, which did not have altered Ca^2+^ sensitivity and did not develop hypertrophy yet displayed diastolic dysfunction^55, 56^. This may be explained by the species differences between the mouse and human, particularly for HCM^57^,and the experimental models used (skinned cardiomyocytes vs hcRTFs). Further studies are needed to dissect these potential differences.

### 4.2 Proteomic and post-translational changes

To further emphasize the molecular changes caused by the *TNNT2* R278C^+/-^ variant, we explored the proteomic profile of both cell lines using mass spectrometry. Most species develop many temporally and spatially controlled muscle myosin heavy chain (MYH) paralogs. This implies that diverse MYH paralog features are required for determining contractility qualities. For example, the myosin α heavy chain (MYH6) has fast twitch mechanics response to excitation-contraction coupling, while β-myosin heavy chain (MYH7) has a slower but more powerful stroke^58^. Therefore, the detected increase in MYH6 expression in our study could be a result of increased contractility and cross bridge interactions with the thin filament because of the increased myofilament calcium sensitivity.

Myosin light chain-2 (MYL2/MLC-2) is a major sarcomeric protein which is also referred to as the regulatory light chain (RLC) of the contractile unit. MYL2 plays an important role in embryonic heart muscle structure and function, its phosphorylation regulates cardiac myosin cycling kinetics, rotation, and contractility^59^. We found a significant decrease in MYL2 expression levels in R278C^+/-^ hiPSC-CMs. This deviation could be attributed to the increased Ca^2+^ sensitivity instigated by the HCM-causing mutations. Hypersensitive myofibrils result in prolonged crossbridge formation, which in turn may result in a disrupted need for regulation by MYL2, leading to a negative response of decreased expression in the variants^60^.

The presence of mechanical strain on the Z-disc may lead to altered transcriptional regulation through mechanosensitive pathways, affecting the expression profile of several downstream proteins, such as MYH^61, 62^. In this study, several alterations were found to be directly or indirectly associated with Z-disc disorganization and myofibrillar disarray. For instance, α-actinin is a major protein found in the Z-disc and is involved in the crosslinking of actin filaments. In HCM, α-actinin is expected to increase due to the greater mechanical stiffness of the CM^42^. Microtubules also play a crucial role in signal transduction and contractility. There is a growing interest in exploring the role of microtubules in affecting diastolic compliance^63^. The upregulation of tubulin β-4B chain in our study could be indicative of elevated tubulin polymerization and stabilization, which is an early pathological adaptive response in cardiomyopathies and heart failure that can induce stiffness which directly affects the diastolic compliance of the cell.

Our results suggest that R278C^+/-^ hiPSC-CMs undergo a switch to a fetal metabolic program, as demonstrated by high expression of glucose genes and lower expression of fatty acid metabolism proteins. The switch to a fetal metabolic profile is commonly seen in HCM and heart failure and may initially be an adaptive response^49^. We additionally found protein changes suggestive of impaired bioenergetics and ATP synthesis or utilization. Impaired sarcomere energetics in HCM may provoke mitochondrial dysfunction, reduced ATP production, and increased reactive oxygen species (ROS) generation. All these indices are directly related to a potential pathogenic decline in the rate of the electron transport chain, leading to lower amounts of respiratory complex proteins^49, 64^. Our findings both coincide with and differ from those found by Gilda e*t al* in a 3-month-old transgenic mouse model in which R278C overexpression caused increased AMP levels indicating metabolic stress yet also decreased glycolysis related proteins^65^. Therefore, the impact of species and age on impaired bioenergetics in HCM needs to be further investigated.

Cardiac contractility is regulated by numerous factors including the phosphorylation of sarcomeric proteins. In relaxed heart muscle there is evidence for a population of myosin cross-bridges with the two heads in an interacting motif (IHM) that establish a so-called super relaxed state (SRX) with low ATPase activity that is sequestered from interacting with the thin filaments. Another population of cross-bridges exists in a disordered relaxed (DRX) state with swaying free heads poised to interact with the thin filaments, hydrolyze ATP, and induce force generation and shortening with calcium activation. In healthy and relaxed human hearts, it is proposed that most myosin heads exist in the SRX state but in pathological conditions like HCM, the IHM state of myosin is disturbed and the percentage of DRX is increased. Phosphorylation of myosin binding protein-C (MyBPC3) and myosin regulatory light chain (MYL2) may promote the DRX state by recruitment of sequestered IHM heads ^51^. Our study provides evidence for the favored transition to the DRX state, in which an increased number of phosphorylated peptides exist in MYBPC3 and MYL2 at sites Ser284 and Ser15, respectively. Sarcomere shortening velocity may also be further affected with the increased phosphorylation of MYBPC3 at Ser284. In a transgenic mouse model, MYBPC3 phosphorylation was found to change both filament orientation and mechanics of contractility by modulation of the SRX state, although no direct effect was observed on Ca^2+^ sensitivity^66^. Additionally, the substitution of arginine (R) with a cysteine (C) in R278C introduces a novel residue for a possible S-glutathionylation. Future studies will explore the modifications on the C278 residue, particularly, S-glutathionylation, which can modulate sarcomeric proteins involved in contraction in relation to changes in the oxidative state of the cell^67^.

### 4.3 Electrophysiological and contractile abnormalities

Tachycardia in HCM patients carrying *TNNT2* variants can be arrhythmogenic. In this regard, the rapid heart rate resulted in atrial fibrillation or sinus tachycardia in over 90% of the arrhythmic episodes in HCM patients who were recipients of implantable cardiac defibrillators (ICDs)^68^. Moreover, sinus tachycardia was observed in 90% of ventricular fibrillation episodes in HCM affected children who were treated with heart rate therapeutics. The results of these studies are consistent with the high incidence of ventricular tachycardia and sudden cardiac arrest possibly linked to high Ca^2+^ sensitivity due to cTnT mutations^5, 12, 17^. Our results in R278C^+/-^ hiPSC-CMs demonstrate that fast stimulating rates can be arrhythmogenic compared to control *TNNT2* hiPSC-CMs. Although the mechanisms of arrhythmogenesis are not clear, alterations in Ca^2+^-handling properties are likely involved. Increased Ca^2+^ sensitivity due to cTnT variants has been previously shown to increase cytosolic Ca^2+^ buffering, since the troponin complex accounts for a substantial portion of cytosolic Ca^2+^ buffering^69, 70^. During diastole, increased Ca^2+^ sensitivity results in the prolongation of Ca^2+^ transient and slower Ca^2+^ decay rate. In previous studies^71, 72^, a theory for the mechanism of intracellular Ca^2+^ alternans was developed. It is called the “3R” theory since Ca^2+^ alternans is promoted by the interactions of random firing of the Ca^2+^ sparks, recruitment of neighboring Ca^2+^ release units (CRUs) for firing (spark-induced sparks), and refractoriness of the CRUs. The recruitment is determined by the coupling strength between CRUs, and Ca^2+^ buffering is one of the parameters affecting recruitment^73^. In this mechanism, Ca^2+^ alternans are promoted by the enhancement of the recruitment^72^. This theory may explain the Ca^2+^ alternans observed in our study. Increased thin filament Ca^2+^ sensitivity appears to be primarily due to the decrease in the Ca^2+^ off rate constant. This reduction/slow-down of the unbinding rate of Ca^2+^ from the myofilament effectively can reduce Ca^2+^ buffering, which can promote Ca^2+^alternans based on the “3R” theory^71–73^.

Moreover, we found the R278C^+/-^ variant to enhance contractility. These findings may be related to the prolonged Ca^2+^ transients seen in R278C^+/-^ hiPSC-CMs. Based on the proteomic results, a reduction in the SRX myosin due to the MYBCP3 and MYL2 phosphorylation may also be regarded as a possible mechanism for increased contractility in R278C^+/-^ hiPSC-CMs^66^. Interestingly, mavacamten was able to alleviate hypercontractility in R278C^+/-^, which may owe to its ability to reduce the number of myosin heads available to interact with actin by promoting the SRX state, which is thought to improve diastolic dysfunction in HCM^74^. Our results are consistent with previous studies examining mavacamten hiPSC-CM models of HCM^74, 75^. Thus, mavacamten may be a promising treatment for thin filament HCM that should be investigated further.

### 4.4 Clinical pathogenicity of *TNNT2* R278C

Like many cardiomyopathy-associated genetic variants, the clinical pathogenicity of the *TNNT2* R278C variant in HCM is controversial. Given that it is now reported in over 50 patients carrying a diagnosis of HCM, it is of critical importance to better understand its role in disease. ClinVar, a publicly accessible online database of human disease associated genetic variants, currently reports conflicting interpretations of pathogenicity^8^. Select laboratories and research groups have reported it to be pathogenic/likely pathogenic, while others have downgraded it to a variant of uncertain significance in HCM (Supplementary Table S1). A major reason to question its association with a relatively uncommon human disease is its higher frequency in the general population (0.06%), as reported in the Genome Aggregation Database (gnomAD)^76^. However, *TNNT2* R278C is generally associated with a late onset of milder HCM, and a relatively higher allele frequency in the general population may be consistent with a low penetrance, high frequency disease-causing variant^77^, as has been described in other forms of genetic cardiomyopathy^78, 79^. Additionally, thin filament (e.g., *TNNT2*) HCM expresses differently than classic thick filament HCM, often causing diastolic heart failure and arrhythmias with borderline ventricular hypertrophy. This makes it more difficult to detect on imaging, particularly in the presence of hypertension and aortic valvular disease, which are prevalent in both the general and late-onset HCM populations (i.e., age 50 or older).

Several studies have reported additional variants in *MYH7* or *MYBPC3* in HCM individuals with the R278C variant. Although this has brought into question whether R278C is in itself pathogenic, it is important to note that HCM cases have been reported without the presence of additional variants, and not all of the additional variants identified are definitively or likely pathogenic. As described in ClinVar (Supplementary Table S1), 14 out of 47 R278C^+/-^ carriers who had HCM possessed an additional variant, which were in MYH7 or MYBPC3. Of these, only 4 out of the 14 possessed a confirmed pathogenic/likely pathogenic secondary variant. Moreover, evidence of co-segregation exists even in families without secondary variants, although expressivity of the R278C variant is incomplete^9, 11, 12, 15^. Therefore, the capability of R278C to result in disease does not seem dependent on the presence of a secondary HCM-associated variant. However, the co-occurrence of HCM-associated variants can affect expressivity and phenotype severity by acting synergistically to accentuate disease manifestation^80^. For instance, Gimeno *et al.* reported that R278C carriers which carried a pathogenic/likely pathogenic secondary variant in MYH7 (p.Asp928Asn) presented with a severe HCM phenotype^81^. Moreover, the presence of an additional variant may not be the only explanation for incomplete expressivity in R278C carriers. The ratio of mutant to wild-type protein expression levels and the impact of post-translational modifications may also contribute. Schuldt et al observed a dose-dependent relationship between the mutant R278C protein expression dose and myofilament function in detergent-extracted cardiomyocytes from the interventricular septum of *TNNT2* R278C^+/-^ patients using troponin exchange experiments. The *TNNT2* R278C^+/-^ variant showed increased Ca^2+^ sensitivity at low doses and a significant increase at intermediate doses^54^. Altogether, our hiPSC-CM study supports the pathogenicity of *TNNT2* R278C^+/-^, although follow up studies are required to address the incomplete penetrance and whether R278C acts as a disease modifier in the presence of secondary variants in thick filament proteins.

### 4.5 Limitations

This study aimed to uncover the proteomic and functional alterations in the early stages of R278C^+/-^ HCM using various approaches. Despite that, several limitations arose that could be improved in future experiments. Firstly, the protein purification protocol is based on a magnetic bead-based solvation layer that captures mostly hydrophilic proteins or soluble portions of proteins. This presents a limitation in studying hydrophobic or membrane-bound proteins (e.g., ion channels). Secondly, functional studies were conducted with only hiPSC-derived cardiomyocytes in 2D monolayers, which could be improved with 3D tissue that includes fibroblasts and endothelial cells, which contribute to the clinical phenotype. Lastly, we did not study the impact of maturation on disease severity, which has been shown to accentuate HCM in hiPSC-CMs studies^82^.

### 4.6 Conclusions

We show that R278C increases Ca^2+^ sensitivity, alters APD and Ca^2+^ transient dynamics, causes hypercontractility, largely disrupts the proteome, and may promote NFAT signaling and a transition towards greater of myosin head in the DRX state. These insights improve our understanding of thin filament HCM and may be transformative for clinical diagnosis and management.

**Table 1:** Average intensity of the slow skeletal (*TNNI1*) and the cardiac (*TNNI3*) troponin 1 paralogs.

| *Average intensity* | *WT* | *R278C* |
| --- | --- | --- |
| *TNNI1* | 32.41 | 31.75 |
| *TNNI3* | 24.69 | 25.34 |
| *TNNI3/TNNI1* | 0.76 | 0.80 |

## Author contributions

SS Experimental: genome editing ; hiPSC differentiation ; optical mapping

AYL Experimental: recombinant protein expression, reconstituted cardiac thin filaments

FJ Experimental: hiPSC differentiation, hiPSC-CM protein extraction, MS proteomics

YM Experimental: hiPSC differentiation; SarcTrack analyses

SD Clinical expertise in familial HCM, ClinVar analysis, manuscript writing

HH Expertise in hiPSC-CM maturation

DHB Expertise in microtubules

TB Experimental: genome editing

BR Experimental: genome editing

SJ Experimental: PDMS matrix design and fabrication

TR Clinical expertise in familial HCM

AC Experimental: Potential energy minimization simulations of the cardiac thin filament

PL Data analysis – thin filament expertise

RJS troponin structure and function expertise

SS Clinical expertise in familial HCM

CT SarcTrack expertise

SL Potential energy minimization simulations of the cardiac thin filament

PL MS proteomics expertise

GFT PI on project, grant holder, project designer and coordinator

## Acknowledgements

The authors are very grateful for the funding supporting this work from several granting agencies. This includes the National Institutes of Health (R01HL137015) to SL, (HLBI RO1 HL158634-01A1) to RJS, Stem Cell Network (FY21 / ACCT2-13) to GFT, and the Canadian Institutes of Health Research (PJT 185981) to GFT. Members of the Cellular and Regenerative Medicine Centre at the BC Children’s Hospital Research Institute are thoroughly indebted to the BC Children’s Hospital Foundation and the Mining for Miracles teams for their incredible infrastructural support.

---

**Table S1.**
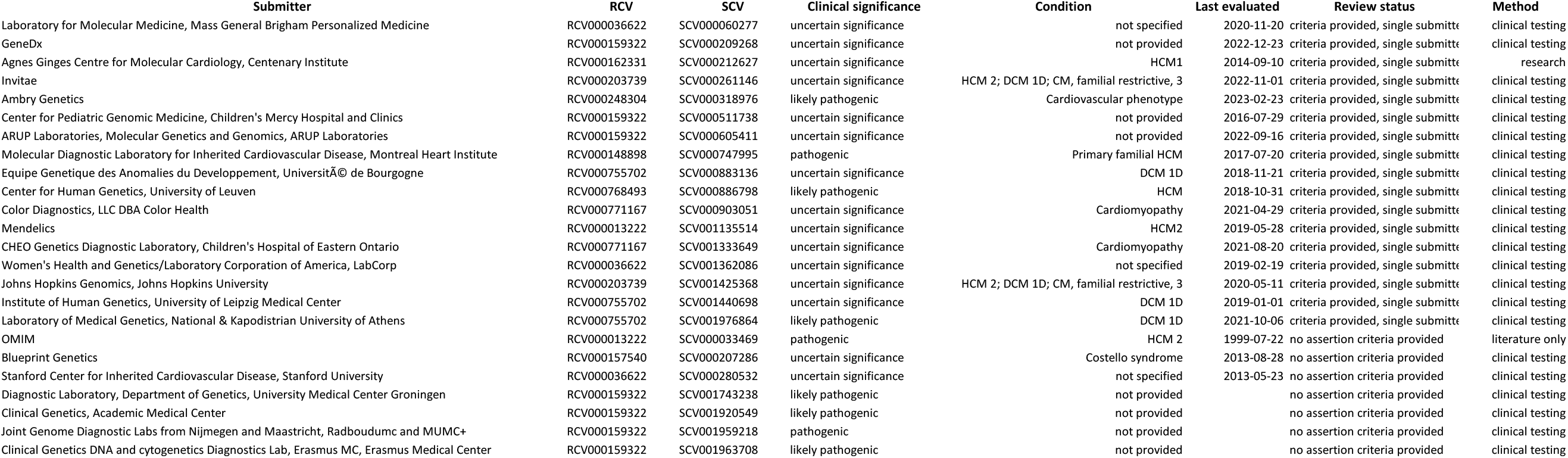
Submissions for variant NM_001276345.2(TNNT2 c.862C-T (p.Arg288Cys)

**Figure S1.**
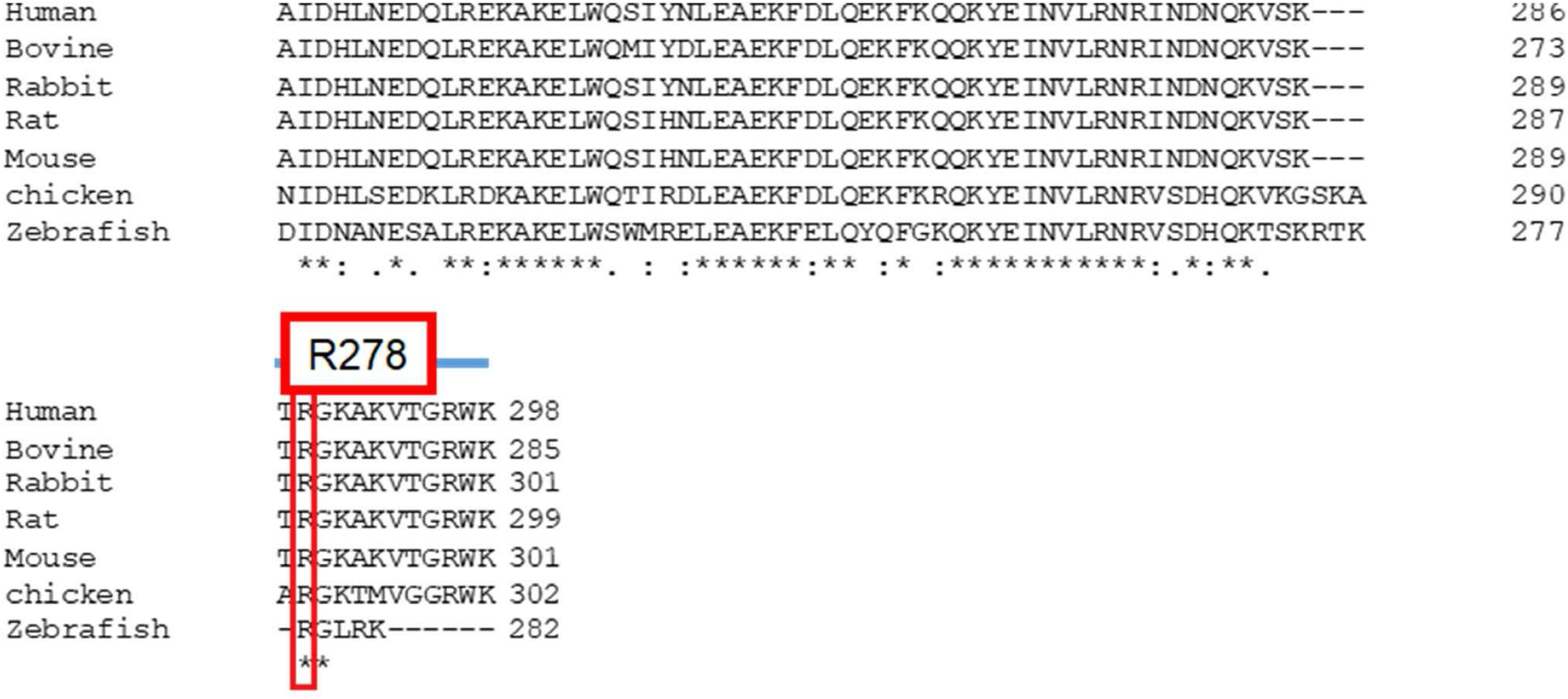
Phylogenetic sequence alignment of cardiac troponin T (TNNT2). A multiple species sequence alignment of the cTnT protein (encoded by *TNNT2*) from human (Uniprot ID: P45379), bovine (Uniprot ID: P13789), rabbit (Uniprot ID: P09741), rat (Uniprot ID: P50753), mouse (Uniprot ID: P50752), chicken (Uniprot ID: P02642), and zebrafish (Uniprot ID: Q90Y46) was performed using Clustal Omega. The equivalent residue R278 among all species are boxed in red and is shown to be highly conserved.

**Figure S2.**
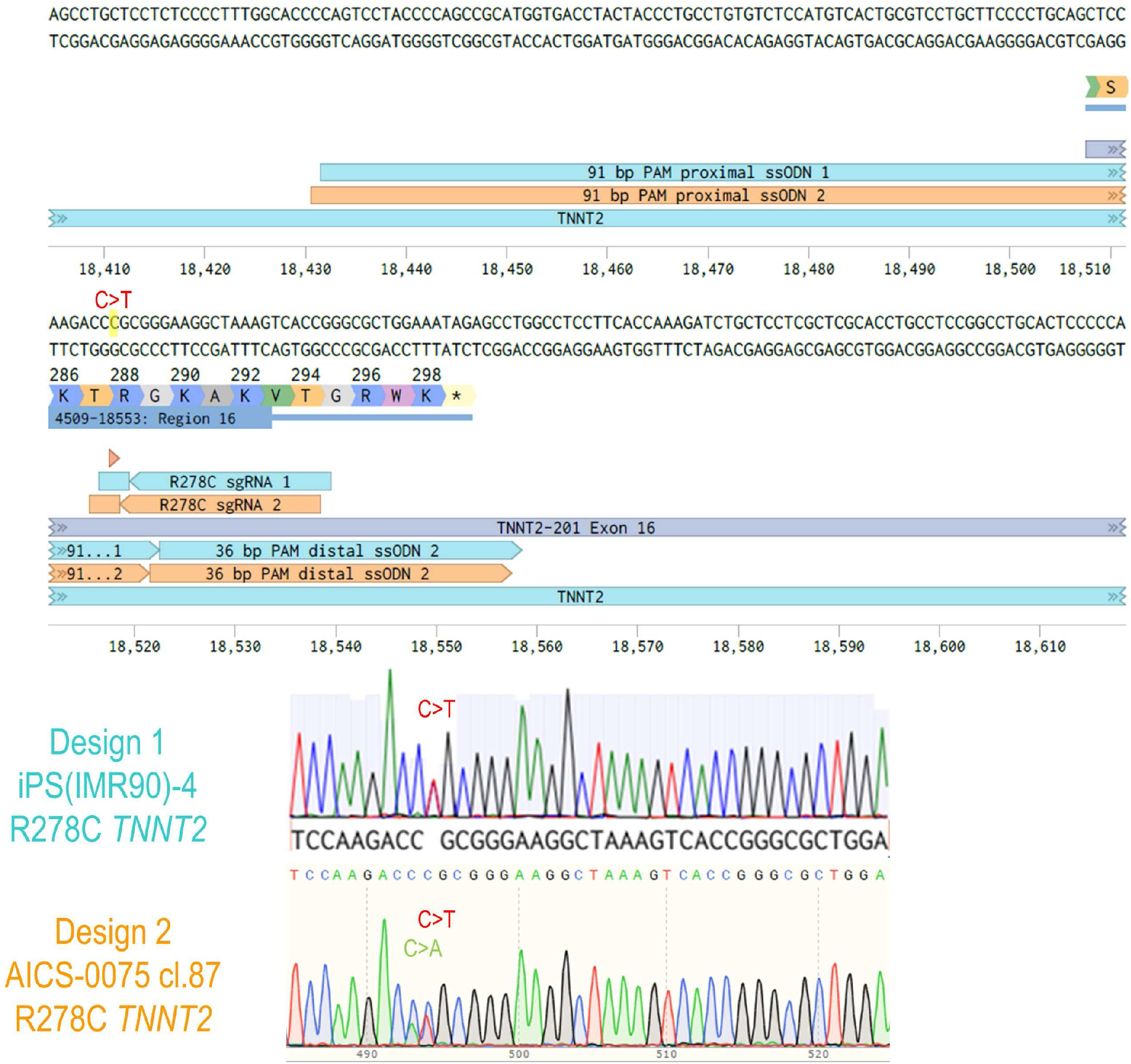
Genome Editing to Generate TNNT2 R278C^+/-^ and WT hiPSC-CMs. Panel A. Schematic of CRISPR/Cas9 design strategies to target R278 in *TNNT2* (R288 prior to postnatal Exon 5 alternative splicing) in two different hiPSC lines iPS(IMR900)-4 (sgRNA 1, ssODN 1, teal) and AICS-0075 cl.87 (sgRNA 2, ssODN 2, orange). The sequence shows the C>T mutation locus for R278C (highlighted in yellow). Panel B. Representative chromatograms of exon 16 in *TNNT2* from isolated iPS(IMR900)-4 (top) and AICS-0075 cl.87 (bottom) lines showing the heterozygous C>T base pair change as double-peaks post-editing for R278C. The C>A silent PAM site mutation is also shown in Design 2. The figure was created using Benchling and SnapGene software. WT = wild-type; hiPSC = human induced pluripotent stem cells; sgRNA = single guide RNA; ssODN = single-stranded oligonucleotide.

